# Archaic introgression and the distribution of shared variation under stabilizing selection

**DOI:** 10.1101/2024.08.20.608876

**Authors:** Aaron P. Ragsdale

## Abstract

Many phenotypic traits are under stabilizing selection, which maintains a population’s mean phenotypic value near some optimum. The dynamics of traits and trait architectures under stabilizing selection have been extensively studied for single populations at steady state. However, natural populations are seldom at steady state and are often structured in some way. Admixture and introgression events may be common, including over human evolutionary history. Because stabilizing selection results in selection against the minor allele at a trait-affecting locus, alleles from the minor parental ancestry will be selected against after admixture. We show that the site-frequency spectrum can be used to model the genetic architecture of such traits, allowing for the study of trait architecture dynamics in complex multi-population settings. We use a simple deterministic two-locus model to predict the reduction of introgressed ancestry around trait-contributing loci. From this and individual-based simulations, we show that introgressed-ancestry deserts are enriched around such loci. When introgression between two diverged populations occurs in both directions, as has been inferred between humans and Neanderthals, the locations of introgressed-ancestry deserts will tend to be shared across populations. We argue that stabilizing selection for shared phenotypic optima may explain recent observations in which regions of depleted human-introgressed ancestry in the Neanderthal genome overlap with Neanderthal-ancestry deserts in humans.

## Introduction

Genomic surveys of natural systems show that historical admixture among diverged populations and closely related taxa commonly occurs (Brandvain *et al*., 2014; Skoglund *et al*., 2015; Suvorov *et al*., 2022) and is widespread in primate (Tung and Barreiro, 2017; SØRENSEN *et al*., 2023) and hominin (Wolf and Akey, 2018; Peter, 2020) evolution. Admixture is therefore a frequent driver of phenotypic and molecular variation and can contribute to the genetic architectures of complex traits. In humans, archaic introgression from Neanderthals and Denisovans has attracted considerable attention, including efforts to describe the historical processes leading to observed distributions of introgressed DNA in present-day populations (PRÜFER *et al*., 2014; Villanea and Schraiber, 2019; Chen *et al*., 2020) and the contribution of introgressed variation to quantitative traits (Sankararaman *et al*., 2016; Wei *et al*., 2023).

Once introduced through admixture, introgressed alleles may be selected for or against. Some introgressed haplotypes were likely positively selected in modern *Homo sapiens* (here, “humans”) (Huerta-SÁNCHEZ *et al*., 2014; Racimo *et al*., 2017; Enard and Petrov, 2018; Gower *et al*., 2021), possibly due to locally adaptive variation that provided fitness advantages as humans encountered novel environments. Despite some cases of adaptive introgression, most introduced functional alleles likely were selected against in humans (Harris and Nielsen, 2016; Juric *et al*., 2016; Veller *et al*., 2023). Since Neanderthal and Denisovan population sizes were relatively small for hundreds of thousands of years, theory predicts they would have accumulated deleterious variation at an increased rate. Introgressed haplotypes loaded with more deleterious mutations would have been rapidly removed by selection after admixture. Mapping the distribution of Neanderthal-introgressed haplotypes in humans shows a reduction of Neanderthal-related ancestry in coding and regulatory regions (Petr *et al*., 2019; Telis *et al*., 2020; Yermakovich *et al*., 2023). These “deserts” of Neanderthal ancestry support the hypothesis that introgressed functional alleles were selected against (Sankararaman *et al*., 2014, 2016).

There is growing genetic evidence that Neanderthals reciprocally received genetic material from early humans (Kuhlwilm *et al*., 2016; Hubisz *et al*., 2020; Harris *et al*., 2023; Li *et al*., 2024). This gene flow occurred tens to hundreds of thousands of years prior to Neanderthal introgression in humans during the global dispersal of modern humans around 60 ka. The earlier admixture is supported by *H. sapiens* outside of African around 200–100 ka (Schwarcz *et al*., 1988; GRÜN *et al*., 2005; Harvati *et al*., 2019; Beyer *et al*., 2021), potentially overlapping with Neanderthals and providing opportunities for early contacts. While estimates of the genomic contribution of early humans to Neanderthals vary, around 6% of later Neanderthal genomes may trace through this admixture event (Harris *et al*., 2023). Under a “load” model, if human-related haplotypes carried fewer deleterious alleles due to their larger long-term effective population size, human-introgressed DNA would have been favored in Neanderthal genomes. The replacement of Neanderthal mitochondrial and Y chromosomes by early human haplotypes appears to support this model of post-admixture positive selection in the Neanderthal lineage (Posth *et al*., 2017; Petr *et al*., 2020).

Models for selection against introgressed alleles are often based on deleterious load or hybrid incompatibilities (Muller, 1942). These models, founded in population genetics theory, rarely take into account selection operating on phenotypic traits or the relationship between genetic and phenotypic variation. Many phenotypic traits are thought to be under stabilizing selection (Sanjak *et al*., 2018; Sella and Barton, 2019), including gene regulation (Gilad *et al*., 2006; Hodgins-Davis *et al*., 2015; Price *et al*., 2022). Because some of the strongest signals of selection against Neanderthal-introduced alleles are in regulatory regions (Sankararaman *et al*., 2014), stabilizing selection on quantitative traits may be particularly relevant to the dynamics of functional genetic variation after introgression among hominins.

Stabilizing selection acts to maintain the phenotypic distribution of a trait near some optimum, which is achieved by reducing phenotypic variation (Figure 1A). When the mean phenotype of the population is close to the phenotypic optimum, classical models predict that the minor allele at a trait-affecting locus is selected against, with allele-frequency dynamics equivalent to underdominant selection (Robertson, 1956). This has proven to be a useful model for understanding genetic architectures of traits under stabilizing selection in single-population settings (e.g., Keightley and Hill, 1988; Simons *et al*., 2018; Hayward and Sella, 2022).

**Figure 1:**
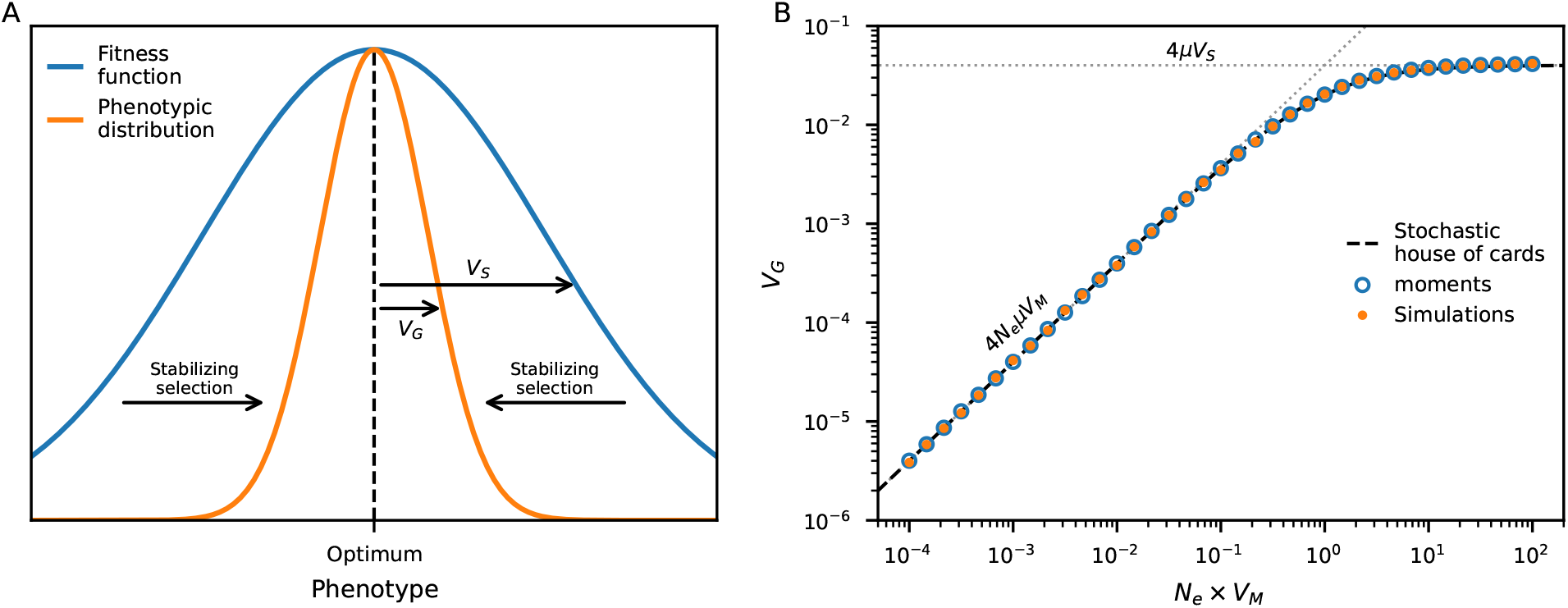
Additive genetic variance under stabilizing selection. (A) Stabilizing selection acts to maintain phenotypic values of individuals in the population near some optimum. Throughout, we assume a Gaussian fitness function. (B) With low mutational variance (*V*_*M*_), the expected additive genetic variance (*V*_*G*_) is proportional to the population-scaled mutation rate. When *V*_*M*_ is large (*N*_*e*_*V*_*M*_ *>* 1) so that mutational effects can be strong, *V*_*G*_ is independent of *V*_*M*_. The stochastic house-of-cards model (Eq. 1) interpolates these regimes, assuming steady-state dynamics (BÜRGER *et al*., 1989). Expected *V*_*G*_ computed using an SFS approach (developed here using `moments`, Jouganous *et al*., 2017) matches simulations assuming linkage equilibrium between loci affecting the trait, which align closely with the stochastic house-of-cards approximation. Here *V*_*S*_ = 1, *N*_*e*_ = 10,000, the mutation rate *μ* = 0.01 (per haploid), and mutation effects are drawn from a normal distribution with mean zero and variance *V*_*M*_.

In diverged populations, the genetic variation contributing to a trait under stabilizing selection has a higher rate of turnover compared to neutral evolution (Yair and Coop, 2022). We therefore expect a rapid divergence of trait architectures in the human and Neanderthal lineages at trait-contributing loci, even when the mean phenotype in each population remains close to the same trait optimum. When a derived allele at high frequency in one population is introduced to another population in which it was previously absent, it will be at low frequency (if the admixture proportion is low) and subsequently selected against. Likewise, if the ancestral allele is reintroduced to a population fixed for a derived allele, the ancestral allele will be at low frequency and will be selected against. Selection acts against the introgressed allele in either case, whether it is ancestral or derived and regardless of the historical relative sizes of the populations involved. In concurrent work, Veller and Simons (2024) demonstrate this effect by deriving the expected decay of minor parental ancestry under stabilizing selection after admixture. This prediction contrasts with the population-genetics load model, in which haplotypes with fewer deleterious variants (such as those from the population with larger historical size) are favored after introgression in either direction.

In this article, we show that admixture between diverged populations results in selection against the minor parental ancestry at loci contributing to a trait under stabilizing selection. We develop a numerical approach based on the site-frequency spectrum to predict genetic variation under complex demographic scenarios, which we use to partition predicted trait heritability by introgressed and non-introgressed variation. Using simulations with linkage, we demonstrate that deserts of introgressed ancestry form around trait-contributing loci. When gene flow occurs bidirectionally, such deserts will tend to overlap in location across populations. We argue that stabilizing selection on shared trait optima may explain the overlap of introgressed-ancestry deserts in human and Neanderthal genomes after reciprocal introgression (Harris *et al*., 2023).

## Model and Methods

We consider a polygenic trait for which an individual’s additive genetic value is the sum over all effects of alleles in their genome. For individual *i, G*_*i*_ = Σ_*l*_ *g*_*l,i*_*a*_*l*_, where *a*_*l*_ is the effect size of the derived allele at locus *l*, and *g*_*l,i*_ ∈ {0, 1, 2} is their genotype at that locus (i.e., the number of derived alleles they carry). With linkage equilibrium between trait-affecting loci, the expected additive genetic variance is 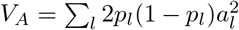, where *p*_*l*_ is the allele frequency at locus *l*. We ignore dominance and epistasis (often argued to be a reasonable modeling choice, e.g., Barton and Keightley (2002); Zhang *et al*. (2002)), so that *V*_*G*_ = *V*_*A*_. We further ignore environmental effects and noise (under the assumption of additive environmental effects, as in Lande, 1975; Simons *et al*., 2018), so that only genetic effects are considered (so that *V*_*P*_ = *V*_*G*_).

Stabilizing selection acts to reduce phenotypic variation around the optimum value *O*, typically set to zero, and we assume a Gaussian fitness function (Figure 1A) so that relative fitness (relative to an individual with optimal phenotype) is given by 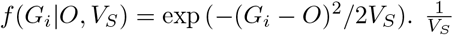 is interpreted as the strength of selection on the trait, so larger *V*_*S*_ implies weaker selection. For a population with mean phenotype at or very close to the optimum, the mean fitness of the population [assuming a normal distribution of phenotypic values in the population, (Turelli and Barton, 1994; Urban *et al*., 2013)] is

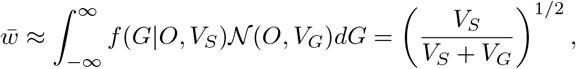

so that as the genetic variance increases, mean fitness decreases.

### Mutation rates, effect sizes and genetic variance

If all alleles contribute equally to the trait with effect sizes ±*a* occurring in equal proportion, Keightley and Hill (1988) showed that the dynamics of *V*_*G*_ can be approximated with the recursion

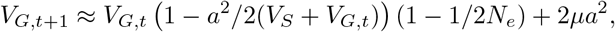

where *μ* is the per-haploid, per-generation rate of mutation. In the large-population-size limit, this gives the well-known result for steady-state additive genetic variance,

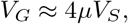

provided *V*_*G*_ ≪ *V*_*S*_.

Mutations will not generally all have the same effect size, but rather are drawn from some distribution. Here, when modeling mutation effects as non-constant, we assume effect sizes follow a normal distribution with mean 0 and variance *V*_*M*_. When population sizes or effect sizes are small, drift dominates the dynamics of *V*_*G*_, and at steady state Lande (1976) found

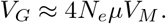

Interpolating between the drift- and selection-dominated regimes,

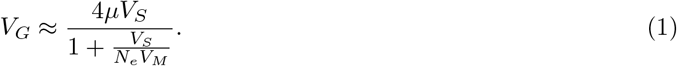

This, the “stochastic house-of-Cards” (SHC) approximation (Figure 1B), was given by BÜRGER *et al*. (1989) and is discussed in detail in Walsh and Lynch (2018, Ch. 28).

### Approximating allelic dynamics via underdominance

As initially shown by Robertson (1956) (see also, Keightley and Hill, 1988; Simons *et al*., 2018), when the mean phenotype of the population is close to the optimal phenotypic value, stabilizing selection results in selection against the minor alleles at loci contributing additively to the trait, with dynamics mirroring symmetric underdominance. In general, for an allele at frequency *p* with selection coefficient *s*, the expected change in *p* over one generation is

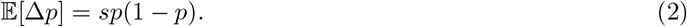

In our case, the selection coefficient *s* depends on the strength of selection on the trait *V*_*S*_ and the effect size *a*, as well as the frequency of the allele, so that

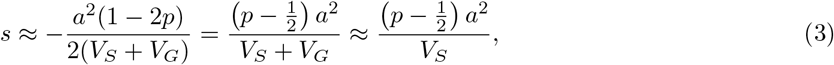

when *V*_*G*_ ≪ *V*_*S*_. This model assumes linkage equilibrium between trait-contributing loci. We note that LD is expected to be generated between loci even in the absence of linkage (Bulmer, 1971), and Negm and Veller (2024) recently showed how to correct for its effect on the selection dynamics of a focal allele.

Because *a*^2^*/*(*V*_*S*_ + *V*_*G*_) is always positive for any *a* ≠ 0, we see that selection pushes allele frequencies to zero if *p <* 1*/*2 and to one if *p >* 1*/*2, resulting in symmetric underdominance. Details are shown in Appendix A.

### Computing expected genetic variance from the SFS

We model allelic dynamics using the diffusion approximation, where the expected change in mean allele frequency per generation is

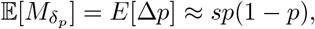

with *s* defined above, and the expected change in variance of the allele frequency per generation is

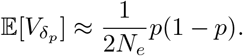

We extended the `moments-`based solution for the sample site-frequency spectrum (SFS) (Jouganous *et al*., 2017) to include underdominance with given selection coefficient *s* (Appendix B). The contribution of alleles with effect size *a* to the total genetic variance *V*_*G*_ is found by computing the expected pairwise diversity from the SFS (with sample size *n*, denoted Φ_*n*_), as

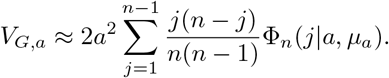

Here, *μ*_*a*_ is the mutation rate of alleles with effect size *a*. Then assuming a normal distribution of effect sizes for new mutations, the total genetic variance is

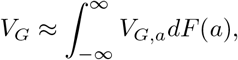

where *F* (*a*) is the cumulative distribution function of 𝒩 (0, *V*_*M*_). *V*_*G*_ can be computed periodically in this way to predict trajectories in non-equilibrium settings (Figure 2).

**Figure 2:**
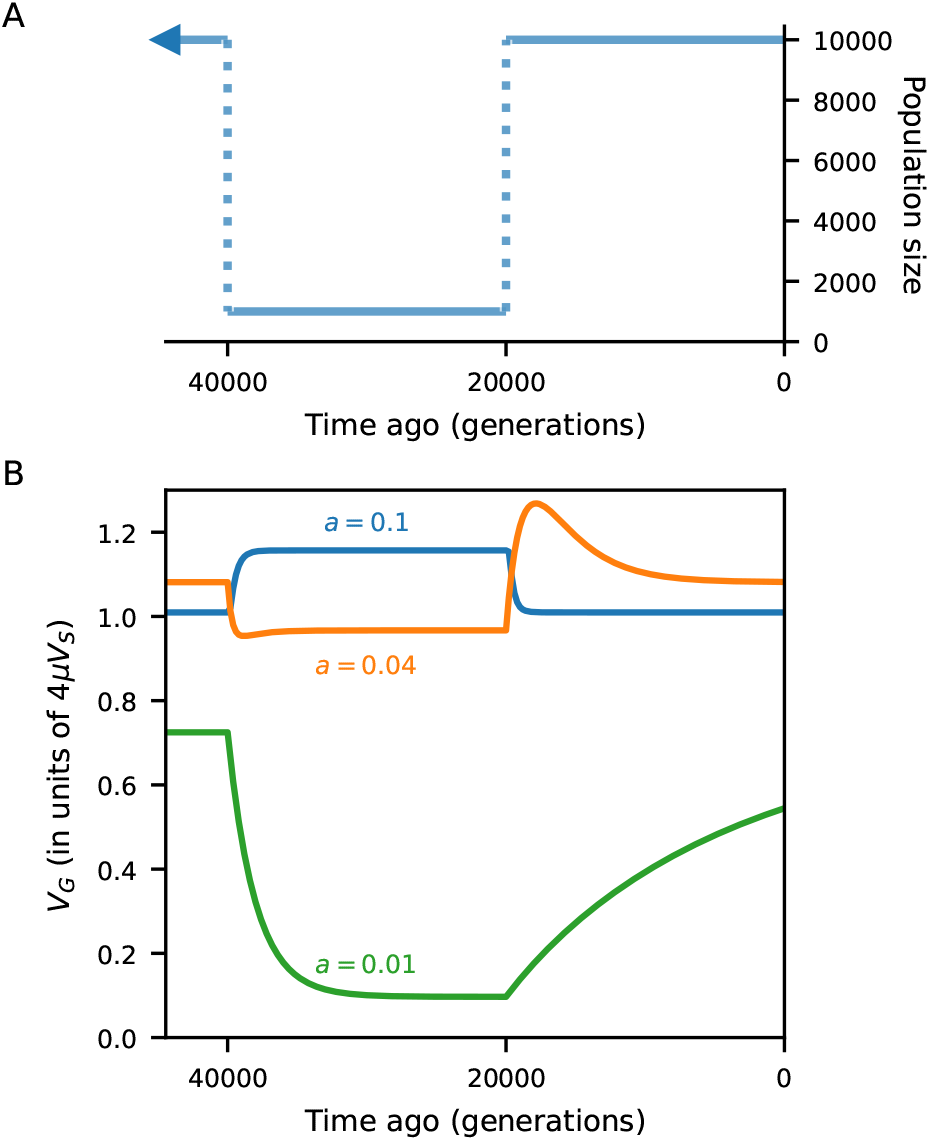
Population size changes and additive genetic variance. We used `moments` to track expected genetic variance of a trait under stabilizing selection as the population changes size. (A) The population goes through a 10-fold size reduction, followed by a recovery. (B) All mutations have effect sizes ±0.01, 0.04, or 0.1, with equal probability of being trait increasing or decreasing. Depending on the selection coefficient (Equation 3) compared to *N*_*e*_, *V*_*G*_ could increase or decrease after a sudden population size change. Predictions using `moments` match simulations with linkage equilibrium between trait-affecting loci (Figures S2–S4).

If *V*_*G*_ is non-negligible compared to *V*_*S*_, ignoring *V*_*G*_ and using *s* = (*p* − 1*/*2)*a*^2^*/V*_*S*_ leads to deflated estimates of additive genetic variance (Figure S1). When mutation rates are large so that *V*_*G*_ is not small compared to *V*_*S*_, using *s* = (*p* − 1*/*2)*a*^2^*/*(*V*_*S*_ + *V*_*G*_) provides estimates of *V*_*G*_ that closely match simulations assuming linkage equilibrium between loci (Figures 3 and S2–S4). Because *V*_*G*_ can vary over time, this means that *s* is no longer constant and can change due to factors such as non-constant demography that increase or reduce *V*_*G*_.

**Figure 3:**
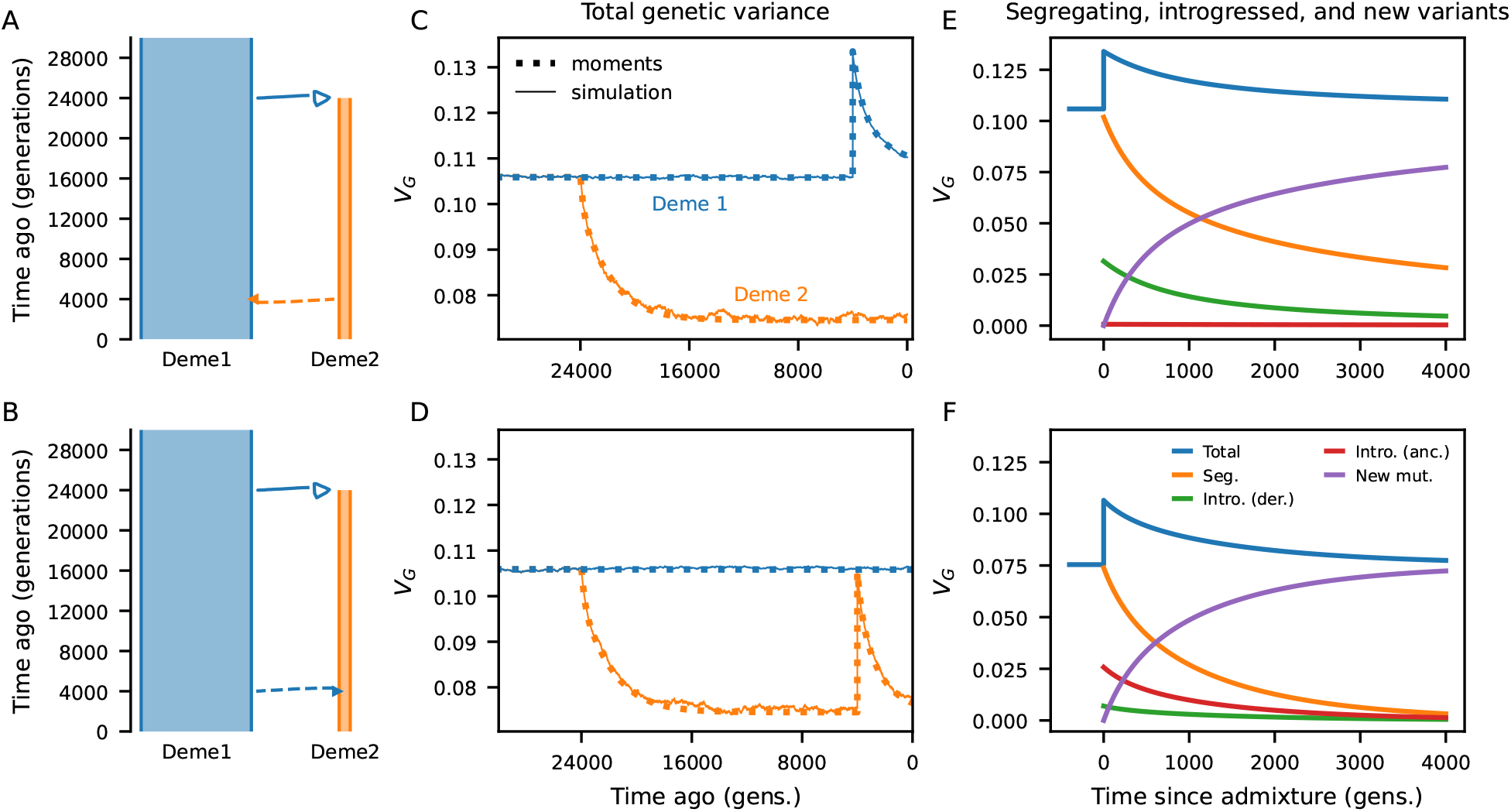
Genetic variance of a trait under stabilizing selection with multi-population demography. (A,B) Two simple demographic models, in which Deme 1 has size 10,000 and Deme 2 has size 1,000. In each scenario, we allow 5% admixture from one deme to the other after begin isolated for 20,000 (2*N*_*e*_) generations. (C,D) We compared predicted (additive) *V*_*G*_ from the site-frequency spectra (using `moments`) to simulations without linkage between trait-affecting loci. In Deme 2, *V*_*G*_ decreases due to their population size reduction. After admixture in both directions, *V*_*G*_ increases in the recipient deme and then decays to steady-state levels. Predictions from `moments` closely match observed *V*_*G*_ in simulations. Here, mutations were drawn from a normal distribution with *V*_*M*_ = 0.0025. Other parameters: *μ* = 0.025, *V*_*S*_ = 1, and the optimal phenotype remained the same in each population. (E,F) Using `moments`, we partitioned additive contributions to *V*_*G*_ by mutations that were previously segregating in the focal population, introgressed variants (either derived or reintroduced ancestral alleles), and by new mutations since the time of admixture. In both demographic scenarios, introgressed variants contribute a substantial proportion to *V*_*G*_, though it is primarily composed of introduced derived alleles when admixture is from the small to the large population, while primarily reintroduced ancestral alleles under the reverse direction of gene flow. In both cases, *V*_*G*_ is increasingly due to new mutations as the genetic architecture of the trait turns over with time since admixture.

### Demographic history

Our numerical solution for the SFS allows for non-constant population size histories, population splits, continuous migration and admixture. Here, we consider relatively simple scenarios involving population splits with subsequent introgression events. We focus on parameter regimes relevant to human-Neanderthal history. In a simple toy model a population of size *N*_*e*_ = 10,000 splits into two, one remaining size 10,000 and the other shrinking to size 1,000. They remain isolated for 2*N*_*e*_ generations (or 500 thousand years, assuming an average generation time of 25 years) and then introgression occurs from one branch to the other, contributing 5% ancestry to the recipient population (Figure 3A,B).

The second model is meant to more closely resemble inferred human-Neanderthal history, in which the ancestral population of size *N*_*e*_ = 10,000 splits at 600 kya into the human and Neanderthal branches, with effective sizes 10,000 and 2,000, respectively. At 250 ka, an early human-to-Neanderthal introgression contributes 5% ancestry to Neanderthals. The human branch shrinks to size 1,000 60 ka, followed by exponential growth to size 20, 000 at present time. Neanderthals contribute 2% ancestry to this bottlenecked and expanding human population at 50 ka, after which they go extinct (Figure 4A).

**Figure 4:**
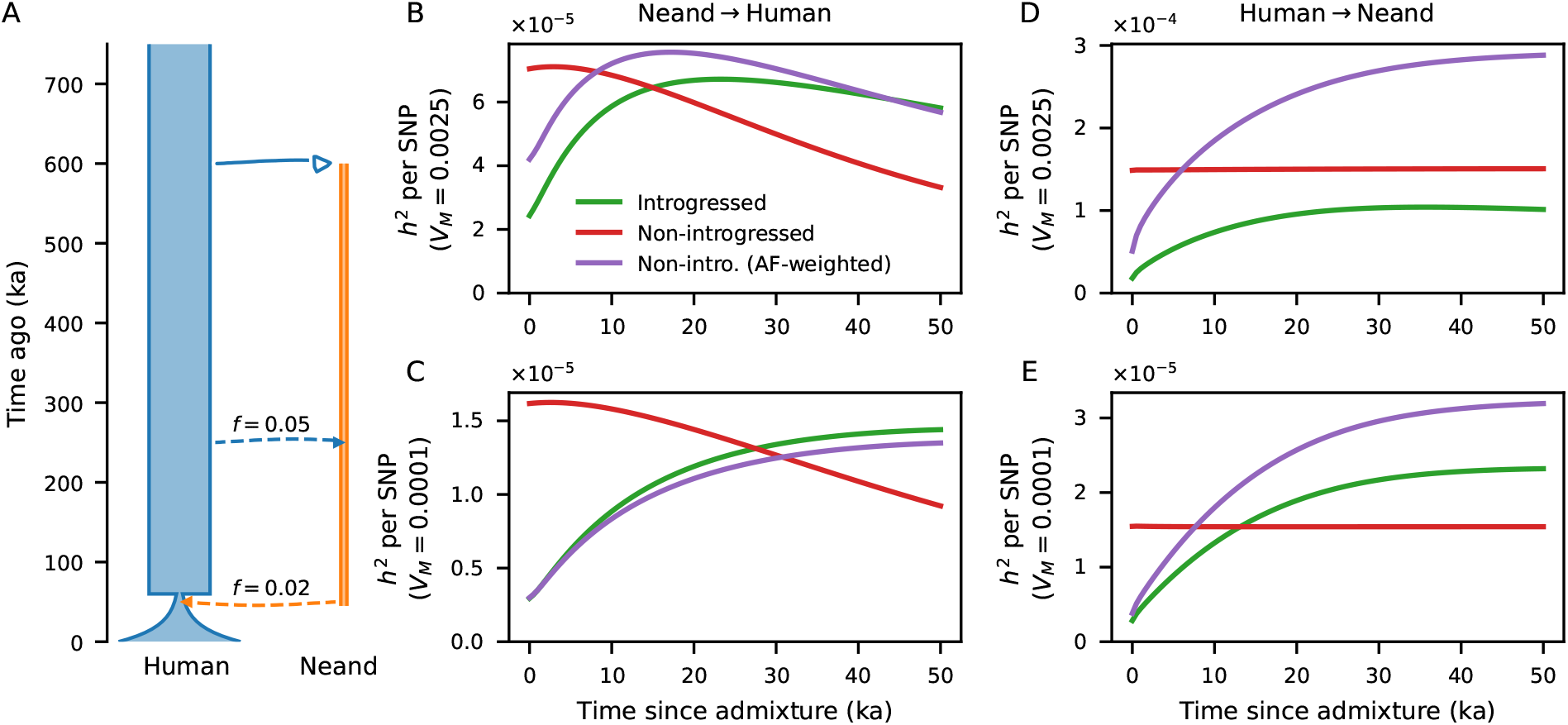
Allele contributions to heritability under human-Neanderthal reciprocal introgression. (A) A model of divergence and admixture between humans and Neanderthals. Using `moments`, we computed predicted *V*_*G*_ over time, partitioned by variation that was introgressed vs. non-introgressed (Figure S7). (B-E) Predicted per-SNP contributions to genetic variance (*h*^2^ per SNP) is plotted over the 50 thousand years following introgression. For non-introgressed variants, we also plot *h*^2^-per-SNP weighted by allele frequencies matching those of introgressed variants. These are shown for (B,C) Neanderthal-to-human introgression 50 kya, (D,E) human-to-Neanderthal introgression 250 kya, (B,D) *V*_*M*_ = 0.0025, and (C,E) *V*_*M*_ = 0.0001.

In each scenario, we tracked phenotypic variance and genetic variation to compare simulations to model predictions. The trait optimum was kept at 0 in all populations (no optimum shift occurred), and the strength of selection *V*_*S*_ = 1 remains constant.

### Simulations with free recombination

We compared our `moments`-based predictions for *V*_*G*_ in non-equilibrium settings to simulations assuming linkage equilibrium as well as individual-based simulations with recombination (next section). For both, mutations occur at rate *μ* per haploid genome copy per generation, with effects drawn from a normal distribution with mean 0 and variance *V*_*M*_.

To simulate allele frequency changes under free recombination (assuming linkage equilibrium between all trait-affecting alleles at all times), for each focal segregating allele we integrated over possible genetic backgrounds contributed by all other segregating alleles. Making the assumption that many alleles contribute to the trait, the variance in genetic backgrounds is normally distributed around the mean genetic value 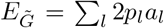 with variance 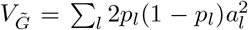, where the sums omit the focal locus. The expected change in frequency was then computed using the approach outlined in Appendix A (without taking the first-order Taylor series approximation for the exponentials). Allele counts in the next generation were then binomially sampled with parameter *p* + 𝔼 [≠*p*] independently for each allele.

### Individual-based simulations with linkage

To include the effects of linkage between multiple selected alleles or selected and neutral alleles, we used `fwdpy11` (Thornton, 2019) to run Wright-Fisher simulations under the demographic models described above. In these simulations, we considered large (1 Morgan, or 100 Mb with a per-base recombination rate of 10^−8^) chromosomes with a uniform recombination landscape. Trait-affecting mutations were either uniformly distributed across the chromosome (Figures 5 and S5–S6) or fell within functional regions (Figures 6 and S8–S12). For the latter, such regions were centered 2 Mb apart and were 100kb in size, so that there were 50 evenly spaced regions across the chromosome. Mutation effect sizes were drawn either as constant values ±*a*, or from a normal distribution with mutational variance *V*_*M*_.

**Figure 5:**
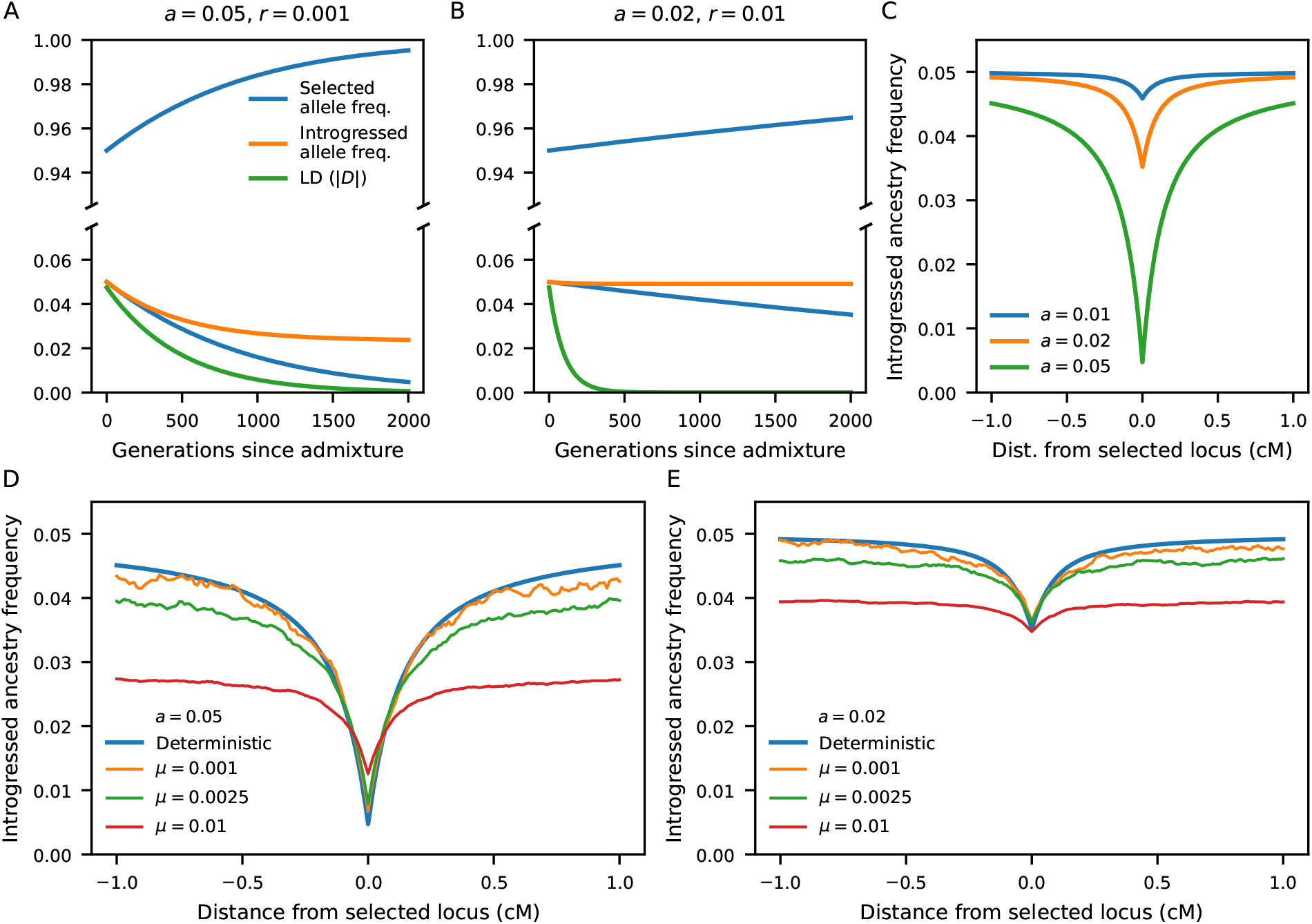
Reduced introgressed ancestry around alleles contributing to a trait under stabilizing selection. Using a deterministic model (Equations 6–8), we model the frequencies of an introgressed trait-affecting allele and a linked neutral allele, initially absent from the recipient population so that their frequencies equal the introgression proportion (*f* = 0.05). The neutral allele frequency is reduced at a rate that depends on the effect size of the selected allele and the probability of recombination between them. LD (as measured by *D* = Cov(*p, q*)) decays to zero over time. (C) Alleles with strong effects are expected to result in a larger depletion of introgressed ancestry around the selected locus. (D,E) Compared to individual-based simulations, the deterministic model predicts the dip in introgressed ancestry around trait-affecting loci, when the mutation rate is low, so that trait-affecting loci are sparsely distributed. When mutation rates (and thus polygenicity) are high, selected alleles tend to be more tightly linked, so that selective interference is more pronounced and local ancestry is affected by multiple selected alleles.

**Figure 6:**
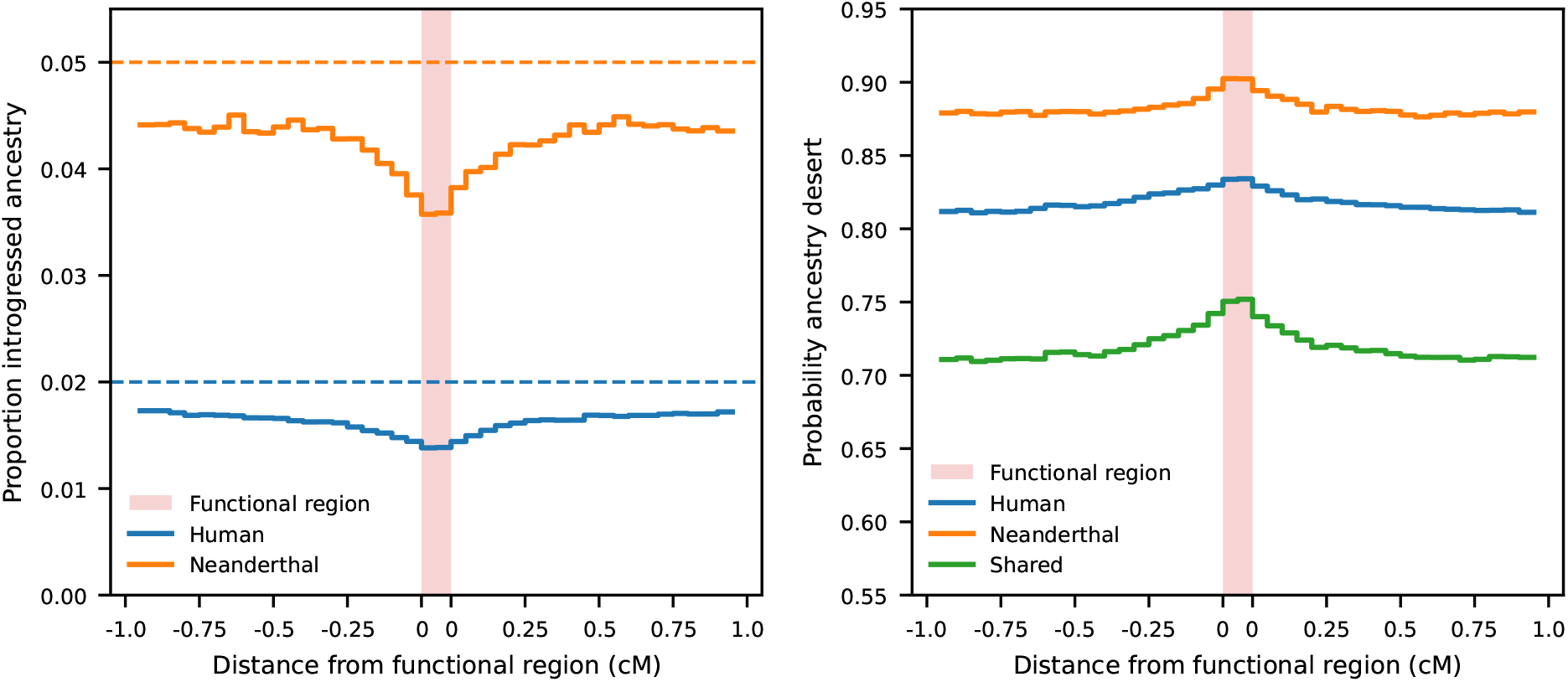
Stabilizing selection causes an enrichment of introgressed ancestry deserts in functional regions. (A) In chromosome-scale simulations under a demography with reciprocal migration between humans and Neanderthals (Figure 4A), stabilizing selection causes a chromosome-wide reduction of introgressed ancestry (below the 5% and 2% introgression proportions). This depletion is most pronounced in “functional” regions that allowed for trait-affecting mutations. (B) Introgressed ancestry deserts are more likely to occur in such functional regions, as are *shared* deserts when compared across samples from humans and Neanderthals. In these simulations, *σ*_*M*_ = 0.02 and *μ* = 0.01. Simulations with different mutational variances are shown in Figures S8–S10. The pattern of shared deserts is not seen in simulations with directional selection (Figures S11 and S12).

In these simulations, we tracked the empirical phenotypic variance (since there is no simulated environmental effect, this is equivalent to *V*_*G*_) in each population each generation. We measured the effects of linked selection on neutral introgressed ancestry using `tskit` to analyze genealogical information (Ralph *et al*., 2020). To measure the reduction in introgressed ancestry around introgressed variants (as shown in Figure 5), we identified each locus with a fixed difference between the parental populations at the time of admixture by preserving the generation immediately preceding admixture. For such a locus, for each sample we determined which (preserved) parental population its ancestry traced to at varying distances from the selected locus. Ancestry proportions were then averaged over each fixed difference. To measure proportions of introgressed ancestry or probabilities of observed ancestry deserts (as in Figure 6), we again used `tskit` to average ancestry proportions of the source population in 50 kb windows. Ancestry deserts were defined as any such window with no ancestry inherited from the source population.

## Data and code availability

All code to run analyses, create figures, and compile this manuscript is available at https://github.com/apragsdale/neanderthal_stabilizing_selection.

## Results

### Additive genetic variance after admixture

We typically expect genetic variance to increase after introgression. The amount that genetic variance increases depends on the allelic differences accumulated between populations and the effects of those alleles. Assuming linkage equilibrium, and ignoring dominance and epistasis, 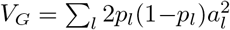. After admixture, with proportion *f* contributed by the population labeled 0 and 1 − *f* by population 1, *p*_*l*_ = *fp*_*l*,0_ +(1 − *f*)*p*_*l*,1_. Plugging into the expression for *V*_*G*_ and after a bit of algebra (Appendix C), we can write the expected genetic variance directly after admixture as

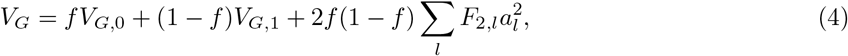

where *F*_2_ = (*p*_0_ − *p*_1_)^2^ is the squared difference in allele frequencies at a locus (Peter, 2016). This result is known (e.g., Tufto, 2000), showing that additive variance is equal to that in the source populations weighted by their contributions, plus a term that depends on the divergence at trait-affecting loci between the populations weighted by the quadratic factor 2*f* (1 − *f*). We note that *V*_*G*_ is expected to increase only in the second generation after admixture, so that selection against introgressed alleles is not immediate (Veller and Simons, 2024). This effect is not captured by this expression.

*F*_2_ at a given locus depends on the demographic history relating the two populations and the effect size at the locus due to selection on the trait. In the infinitesimal limit, involving many loci each of vanishingly small effect, dynamics at a given locus will be approximately neutral, so that *F*_2_ depends only on demographic history. In this case,

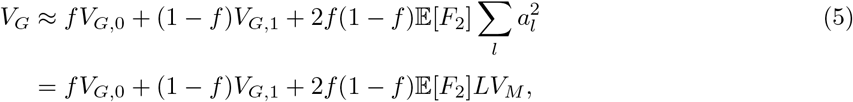

where *L* is the number of trait affecting loci.

### Predicted *V*_*G*_ from the SFS

Expected allele frequency differences (*F*_2_) for selected alleles differ from neutrality. For negative and underdominant selection, *F*_2_ is reduced relative to neutral expectations (Figure S13). Because analytic solutions are unavailable for arbitrary evolutionary scenarios involving multiple populations, we numerically solve for the expected joint distribution of allele frequencies (the SFS) before and after admixture. This provides a numerical solution for expected *V*_*G*_, which can be tracked over time (Methods). Comparing to simulations with free recombination between loci, we observe close agreement with average *V*_*G*_ at all times (Figure 3C,D). In the scenarios tested here, admixture causes a sudden increase in *V*_*G*_ followed by a fairly rapid return to pre-admixture levels, which is recovered by our numerical approach.

Modeling the dynamics of genetic variance using the SFS lets us examine contributions to *V*_*G*_ from different classes of mutations, such as those at different frequencies or arising at different times, and how those contributions change over time. In particular, we may quantify the contribution to *V*_*G*_ from alleles that were already segregating in the recipient population, those that were introduced through introgression, and new mutations since the time of admixture (Figure 3E,F). In many scenarios of interest, in which populations are considerably diverged at the time of admixture, genetic architectures will be largely unique in each population. After mixing, previously segregating and introgressed alleles each contribute to *V*_*G*_ before going to fixation or loss, and the variance of the trait is increasingly due to new mutations.

Introgressed variation can initially make up a considerable portion of *V*_*G*_, with those alleles being either newly introduced derived alleles or reintroduced ancestral alleles. The relative sizes of the two populations impact the numbers of each, as derived alleles will accumulate more readily in a population with smaller effective size. Nonetheless, the overall increase in *V*_*G*_ is similar in both directions of introgression, as the symmetric term 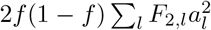 contributes in either case and can be much larger than *f V*_*G*_ from the source population (Equation 5).

### Complex demography and partitioning heritability by origin of alleles

We used a historical model resembling inferred human-Neanderthal history (Figure 4A) to explore the effects of population size changes and reciprocal admixture on the additive genetic architecture of traits under stabilizing selection. As expected (Figure 3), population contractions decrease *V*_*G*_ as drift removes allelic diversity at trait-affecting loci, while introgression increases *V*_*G*_.

Because the genetic architectures considered here are purely additive, we can track mutations in an admixed populations by whether they were previously segregating, fixed or lost in either parental populations or if they arose as new mutations since the time of admixture. Partitioning *V*_*G*_ by contributions from these different sets of mutations (Figure S7), we find it is still primarily contributed by previously segregating, non-introgressed mutations. *V*_*G*_ due to existing mutations decays monotonically over time and is rapidly replaced by new mutations.

The average contribution of introgressed vs. non-introgressed SNPs to *V*_*G*_ (i.e., *h*^2^-per-SNP) can similarly by tracked over time. For the human-Neanderthal demographic model and genetic architectures considered here, the contribution per-SNP of introgressed variants is initially lower than that of non-introgressed variants. These contributions change over time, depending on mutational variance, as well as demography (Figure 4). When weighting *h*^2^-per-SNP of non-introgressed SNPs by matching to allele frequencies of introgressed variants, relative contributions depend sensitively on evolutionary parameters and the time since admixture.

### The effects of linkage

In the preceding sections, we found that approximating the dynamics of trait-affecting alleles using an underdominant selection model (Robertson, 1956) provides an excellent approximation of *V*_*G*_ in complex demographic scenarios. However, this relies on linkage equilibrium between trait-affecting alleles. The inclusion of linkage can cause noticeable distortions of expected *V*_*G*_, so that Equation 1 differs from observed *V*_*G*_ at steady state (BÜRGER *et al*., 1989; BÜRGER and Lande, 1994; Walsh and Lynch, 2018).

To investigate the effects of linkage, we used chromosome-scale individual-based simulations (Thornton, 2019). By varying the mutation rate and the variance of effect sizes of new mutations, we included scenarios ranging from low to high polygenicity and from weak to strong selection on individual alleles. In this and the following sections, we highlight two effects of linkage. First, we observe deviations of *V*_*G*_ from expectations under free recombination, which can be large for highly polygenic traits. Second, selection on introgressed trait-affecting alleles results in a reduction of introgressed ancestry in surrounding regions.

With linkage between two or more loci contributing to a trait under stabilizing selection, linkage disequilibrium (LD) can develop between alleles. Notably, stabilizing selection will lead to coupling LD between mutations of opposite-signed effects and repulsion LD between those of same-signed effects (Bulmer, 1971). This is expected to decrease *V*_*G*_. For lower mutational variances (2*N*_*e*_*V*_*M*_ ≈ 1, Figure S5), we observe such a reduction in *V*_*G*_. With low mutational input, and thus low polygenicity, *V*_*G*_ in individual-based simulations closely matches expectations under linkage equilibrium. With increasing mutation rate, *V*_*G*_ is reduced relative to those expectations. However, when the mutational variance is much larger (2*N*_*e*_*V*_*M*_ ≈ 50), we see the reverse trend (Figure S6). At low mutation rate, there is a close match between observed *V*_*G*_ and expectations, although simulated values are slightly higher. As the mutation rate increases, *V*_*G*_ increases to be relatively much larger than expectations, rather than smaller.

The strength of the deviation of *V*_*G*_ between models with and without linkage depends on a number of factors. The total mutation rate affects not only the polygenicity, with higher mutation rates producing more segregating alleles, but also influences the mean recombination rate between alleles, as they will be more densely distributed in the genome. The distribution of effect sizes plays an important role, as seen by the opposite trends in *V*_*G*_ distortions (Figure S5 vs. Figure S6), which are most apparent with high mutation rates and thus high polygenicity. In these cases, there appear to be complex dynamics involving the reduction in *V*_*G*_ due to the Bulmer Effect (Bulmer, 1971) and an increase in *V*_*G*_ due to selective interference (Hill and Robertson, 1966).

### Introgressed ancestry is reduced around introduced trait-affecting alleles

Because introgressed trait-affecting alleles are selected against as the minor allele, introgressed ancestry segments in the regions surrounding the selected alleles will also be removed due to linkage. The expected reduction in introgressed ancestry will depend on the effect size of the linked trait-affecting allele and the local probability of recombination. We first consider a deterministic model for the frequency dynamics of introgressed alleles (one selected, one neutral) with variable rate of recombination. This simple model ignores the effects of drift and of interference between multiple selected alleles.

We model a trait-affecting locus with a fixed difference between the two parental populations, in which the derived allele may be fixed in either population. At the functional locus, with admixture proportion *f* from the minor parental source, the initial frequency of the derived allele is either *p*_0_ = *f* or *p*_0_ = 1 − *f*. Over one generation, the expected allele frequency at the selected locus is, to leading order in *s*,

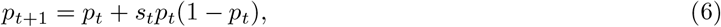

with *s*_*t*_ = (*p*_*t*_ − 1*/*2)*a*^2^*/V*_*S*_ for stabilizing selection (Equation 3).

We consider a neutral locus separated from the selected locus by recombination fraction *r*. Initially, the expected frequency of linked introgressed ancestry is *q*_0_ = *f*, which changes over time due to linked selection on the trait-affecting allele. Letting *D* = Cov(*p, q*) be the standard covariance measure of LD between the alleles at the two loci, *q* is expected to change as

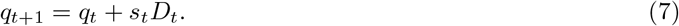

Initially, *D*_0_ = ±*f* (1 − *f*), with *D* being positive if *p*_0_ = *f* and negative if *p*_0_ = 1 − *f*. LD between the loci changes deterministically over time due to both selection and recombination, so that

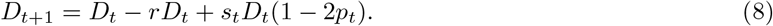

Together, this forms a simple nonlinear system of equations for the deterministic change in allele frequencies at the two loci and LD between them.

Using this model, we predicted the changes in introgressed allele frequencies and LD after admixture for given effect size *a* and recombination rate *r* (Figure 5A,B). As expected, smaller effect sizes result in a slower decay in introgressed ancestry frequency at both selected and linked loci, and larger recombination rates more quickly decouple the linked ancestry from the selected allele dynamics. Thus, the expected reduction in introgressed ancestry is largest for larger effect sizes (Figure 5C), and LD between the selected and linked neutral alleles is largest for neutral sites closest to the selected allele (Figure S14).

We assessed the accuracy of the deterministic two-locus model using individual-based simulations (Thornton, 2019) under a simple demographic model (Figure 3A), with introgression fraction *f* = 0.05. We simulated a single chromosome of length 1 Morgan, with all mutations having effect sizes ±*a* (*a* = 0.02 or 0.05, in separate simulations), and we varied the total per-chromosome mutation rate (*μ* = 0.001, 0.0025 and 0.01). For each fixed-difference mutation in the parental populations, we determined the average introgressed ancestry surrounding such loci 4,000 generations after admixture (Methods).

The deterministic model (Equations 6–8) provides a very good approximation when mutation rates are small (Figure 5D,E). As the mutation rate increases in these simulations, trait-affecting alleles are more densely distributed along the chromosome. Deviations from the deterministic model are due to multiple selected alleles affecting local introgressed ancestry. For the highest mutation rate shown here, there can be many other trait-affecting alleles within a 1 cM window around any focal SNP, distorting dynamics away from predictions under the two-locus model.

### Introgressed ancestry deserts are shared under stabilizing selection and reciprocal introgression

As shown in the previous section, introgressed ancestry is reduced around loci with trait-affecting alleles regardless of the parental population the derived allele is present in. We should therefore expect to observe reductions in introgressed ancestry at the same trait-affecting loci after gene flow in either direction. If introgression occurs in both directions, regions of reduced introgressed ancestry will coincide around such loci, and introgression “deserts” that appear due to selection against trait-affecting alleles will tend to be shared.

To demonstrate this effect, we performed chromosome-scale simulations under a model of reciprocal introgression between humans and Neanderthals (Figure 4A, Harris *et al*., 2023). Mutations affecting a trait under stabilizing selection in each population occurred in 100 kb “functional” regions, spaced 2 Mb apart (Methods). Sampling individuals from both the human and Neanderthal lineages, we observed the average introgressed ancestry proportions were lowest within and surrounding functional regions (Figure 6A). This corresponded to an increased proportion of ancestry deserts within the functional regions (Figure 6B). Functional regions also displayed an enrichment of *shared* ancestry deserts, with the probability of observing ancestry deserts (either within a population or shared) decaying to background levels as the distance from the functional region increases.

## Discussion

Many phenotypic traits are under stabilizing selection (Hodgins-Davis *et al*., 2015; Sanjak *et al*., 2018; Sella and Barton, 2019). This has motivated using stabilizing selection around a shared optimum as a null model for the dynamics of alleles affecting polygenic traits, including in multi-population settings (Yair and Coop, 2022). Stabilizing selection on a trait results in selection against the minor allele, mirroring symmetric underdominance (Robertson, 1956; Keightley and Hill, 1988). Some theoretical and simulation studies have considered the effect of population differentiation or migration on the genetic architecture of a trait under stabilizing selection (e.g., Tufto, 2000; Yeaman and Whitlock, 2011; Yair and Coop, 2022), but most previous work has focused on single population scenarios, often assuming steady state dynamics. Episodes of admixture and introgression commonly occur in many species’ evolutionary histories, so understanding their effects on the architecture of complex traits is needed.

We find a multi-population, non-equilibrium approach for the site-frequency spectrum with underdominance can be used to model the additive genetic variance of a trait under stabilizing selection. While this approach ignores some biological relevancies, such as linkage between sites, non-additivity and pleiotropy, it still provides important insights into the dynamics of trait architectures. For example, we can readily decompose contributions from alleles of different origins, either by source population or mutation time. It may therefore be useful for understanding the contributions of introgressed variants to complex traits, such as hominin-introgressed mutations in humans (Reilly *et al*., 2022; Wei *et al*., 2023). In the limited scenarios explored here, we find that the expected contributions of introgressed variants to heritability are complicated even in the purely additive case, depending on the distribution of effect sizes, demographic history and the time since admixture (Figures 4 and S7).

After admixture between diverged populations, the additive genetic variance of a trait rapidly increases. The genetic variance after admixture depends on both the existing genetic variances within the parental population and their divergence at trait-affecting loci, measured by *F*_2_ (Equation 5) (see also, Veller and Simons, 2024). *V*_*G*_ is then expected to decay fairly quickly to background levels. During this process, introgressed and pre-existing trait-affecting alleles are replaced by new mutations, turning over the genetic architecture of the trait (Figure 3). Selection occurs against the minor alleles at trait-affecting loci, so that introgressed alleles from the minor parental ancestry, whether derived or reintroduced ancestral alleles, are selected against (Figure 5).

Selection against introgressed alleles also removes linked introgressed ancestry in the surrounding regions. Thus, introgressed ancestry deserts are more likely to form at and around loci contributing to selected traits (Figure 6). This process is symmetric, so that deserts tend to form in the same regions under reciprocal introgression. When comparing distributions of introgressed haplotypes in two diverged populations with recent bi-directional gene flow, we should expect to see an overlap of deserts in genomic regions contributing to traits under stabilizing selection.

The expected pattern of shared ancestry deserts under stabilizing selection differs from models of both deleterious load and incompatibilities, providing testable hypotheses. Load-based models predict that haplotypes that have accumulated more deleterious mutations, e.g., from a population with small long-term effective population size, will be selected against under either direction of gene flow. Introgressed ancestry at a given selected locus will decrease in frequency in one introgression scenario and increase in the other. This may explain the replacement of MT and Y chromosome DNA in Neanderthals by human haplotypes after early human-to-Neanderthal introgression and the absence of such Neanderthal haplotypes in modern humans (Posth *et al*., 2017; Petr *et al*., 2020), but does not broadly match observations across the autosomal genome (Harris *et al*., 2023).

The classical model of Bateson-Dobzhansky-Muller incompatibilities (BDMIs) (Bateson, 1909; Dobzhan-sky, 1936; Muller, 1942) explains the accumulation of reproductive isolation through negative epistatic interactions that are exposed in hybrids. Muller (1942) hypothesized that such BDMIs should most often form between distant or unlinked loci, instead of within a single locus or tightly linked loci. Because theory and experiments show that hybrid incompatibilities are resolved via selection against the minor parental ancestry (Matute *et al*., 2020; Moran *et al*., 2021), ancestry deserts should form in different genomic regions, since different incompatibility alleles are selected against depending on the parental ancestry proportions. While both BDMI and stabilizing selection models predict selection against introgressed alleles, an important distinction is that deserts due to selection against incompatibility loci are not expected to overlap under bidirectional gene flow. While there is little empirical data on the distribution of BDMIs, studies point to interacting BDMI alleles being unlinked (Presgraves, 2003; Li *et al*., 2022) and an asymmetry in the alleles under selection in different introgression scenarios (Maheshwari and Barbash, 2011; Moran *et al*., 2021).

Recently, Harris *et al*. (2023) observed that regions of depleted Neanderthal ancestry in humans overlap more than would be expected by chance with regions lacking human-introgressed alleles in the Altai Neanderthal. Human-introgressed ancestry in the Neanderthal genome is also depleted in functional regions (Harris *et al*., 2023), as is well-documented in humans (Sankararaman *et al*., 2014, 2016). Harris *et al*. (2023) propose that epistatic interactions between introgressed alleles and the recipient backgrounds could drive these patterns, which they interpret as evidence for the initial process of speciation between humans and Neanderthals. However, the observation of shared ancestry deserts does not match expectations under a classic model of BDMIs as described above. Instead, at least some of the pattern may be due to stabilizing selection acting on complex traits, such as gene regulation. Importantly, such overlapping ancestry deserts are expected even when a trait is under stabilizing selection for the same phenotypic optimal value. While the underlying causes of selection on introgressed alleles in humans and Neanderthals remain largely unknown, stabilizing selection provides a well-grounded explanation for observed patterns that should be considered when testing for epistasis, incompatibilities and adaptive introgression.

## Acknowledgments

I am grateful to Kevin Thornton, Bret Payseur, Carl Veller and Yuval Simons for valuable discussions, and Carl Veller for helpful comments on an earlier version of this manuscript. Support for this research was provided by the Office of the Vice Chancellor for Research and Graduate Education at the University of Wisconsin–Madison with funding from the Wisconsin Alumni Research Foundation.

## Appendix

### A Stabilizing selection results in selection on trait-affecting alleles equivalent to symmetric underdominance

This result is well known and has been derived many times before. We include it here for completeness. We consider a polygenic trait under stabilizing selection, and we assume the mean phenotype of the population is close to the optimal value (0), which will be the case if the optimum has not shifted recently. Because many segregating loci are assumed to contribute to the phenotypic variance of the trait, the phenotypic distribution is well-approximated as normally distributed around the optimum: 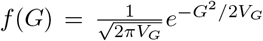. We assume a Gaussian fitness function, so that the fitness of an individual with genotypic value *G* is given by 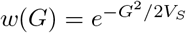.

For a derived allele (labeled 1) with frequency *p*, the expected change in frequency over one generation due to selection (in general) is

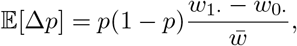

where 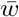 is the mean fitness of the population, and marginal fitnesses for the derived and ancestral alleles are

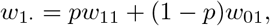

and

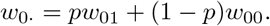

Here, *w*_11_, *w*_01_, and *w*_00_ are the relative fitnesses of individuals who are homozygous for the derived allele, heterozygous, or homozygous for the ancestral allele, respectively.

We partition an individual’s genetic values into contributions from their genetic background 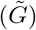 and the focal locus: 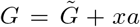, where *x* ∈ 0, 1, 2. We then integrate over genetic backgrounds to find the expected fitnesses of each genotype. As described in SI section 2.1 of Simons *et al*. (2018), because each individual locus contributes a small amount to 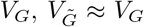 and the fitness function is well approximated as 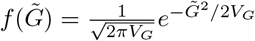. Then, mean fitness

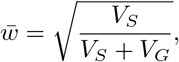

as shown in the main text, and

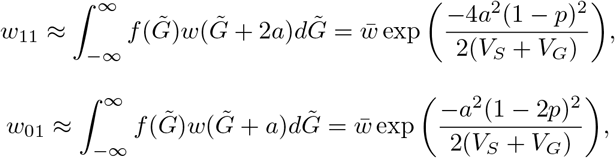

and

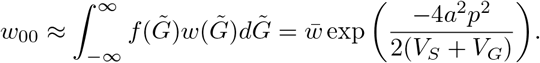

Taking the first-order Taylor series expansion of the exponentials (*e*^−*x*^ ≈ 1−*x*, for small *x*) and combining terms,

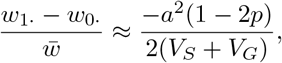

demonstrating the result.

This same result can be found by directly considering the marginal fitness of a focal allele against the expected haploid background, as pointed out by Negm and Veller (2024). Thus, while the selection coefficient takes an equivalent form to underdominance, we note that the mechanism differs: selection acts against the minor allele, rather than against the heterozygous genotype.

### B Moment equations for over- and underdominance

For symmetric underdominance, the relative fitnesses of genotypes *aa* : *Aa* : *AA* are 1 : 1 − *t* : 1. We consider any value of *t* (positive or negative, corresponding to either under- or overdominance); with stabilizing selection, 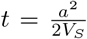, so that heterozygous individuals always have reduced fitness compared to either homozygote.

We extended `moments` (Jouganous *et al*., 2017) to compute the sample site-frequency spectrum Φ_**n**_ for one or more (up to five) populations with sample sizes **n**. This provides a good approximation for the distribution of trait-affecting allele frequencies across multiple populations if the trait optimum is shared across populations and each population’s mean phenotype remains close to that optimum. This is expected to be the case if there are no optimum shifts in any lineage. Hardy-Weinberg equilibrium is assumed at all loci. Accounting for optimum shifts in one or more lineages would require a combination of direct and underdominant selection (e.g., Hayward and Sella, 2022), which we leave for future work.

We refer readers to Jouganous *et al*. (2017) for a detailed introduction to the general moments-based framework for the dynamics of Φ_**n**_. Here, we describe how underdominant selection changes Φ over a single generation. With symmetric underdominance, selection acts on heterozygotes. This can be formulated as some proportion of heterozygotes failing to reproduce in a given generation, with those selected lineages replaced by copies drawn from the full population.

In a single population, we consider *n* tracked lineages of which *i* of those carry the derived allele (Φ_*n*_(*i*) is thus the count of loci with *i* observed derived alleles in a haploid sample of size of *n*). *i* can increase or decrease due to selection “events”. Here, we assume *t* is small enough so that at most a single selection event occurs among the *n* lineages in any given generation. This is a reasonable approximation as long as *t* is not extremely large (Jouganous *et al*., 2017; Krukov and Gravel, 2021) – for selection coefficients induced by stabilizing selection, this is typically a safe assumption.

Two selective events can change *i* to *i* + 1 or *i* − 1: (a) a tracked copy carrying a derived allele is heterozygous (paired with an ancestral allele-carrying copy), selected against, and then replaced by an ancestral allele drawn from the rest of the population, so that *i* → *i* − 1, or (b) a tracked copy carrying the ancestral allele is paired with a derived allele-carrying copy, selected against, and then replaced by a derived allele, so that *i* → *i* + 1. In both cases, we require drawing two additional lineages to find the Φ_*n*_ in the next generation, one for the diploid pair and one for the replacement allele (i.e., 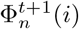 requires 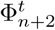). This results in an unclosed system of equations, and we use a quadratic jackknife approximation to approximate Φ_*n*+2_ from Φ_*n*_, as described in Jouganous *et al*. (2017).

The expected increase and decrease of Φ_*n*_(*i*) due to selection is found by enumerating the sampling probabilities of each selective event. For case (a), *i* → *i* − 1 (i.e., Φ_*n*_(*i*) is reduced) with probability

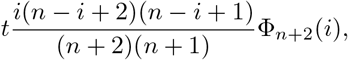

and *i* + 1 → *i* (Φ_*n*_(*i*) increases) with probability

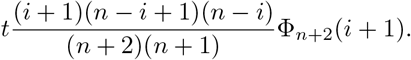

For case (b), *i* → *i* + 1 with probability

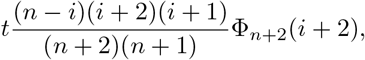

and *i* − 1 → *i* with probability

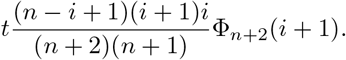

These can be combined (with negative rates for the reduction of Φ_*n*_(*i*)) to describe the change of Φ_*n*_(*i*) for all 0 ≤ *i* ≤ *n*. Figures S15–S17 show that this approach, implemented in `moments`, is accurate compared to discrete Wright-Fisher simulations.

### C Additive genetic variance after admixture

Here, we derive the expected genetic variance after admixture between two source populations (Equation 5). Suppose two population (labeled 0 and 1) diverged some time in the past and then admix in proportions *f* and 1 − *f*.

Assuming random mating, linkage equilibrium, and no dominance or epistasis (so *V*_*G*_ = *V*_*A*_),

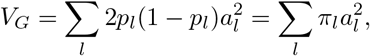

where *π* denotes expected pairwise diversity. At a given locus (dropping the *l*), after admixture the allele frequency is

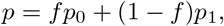

so that

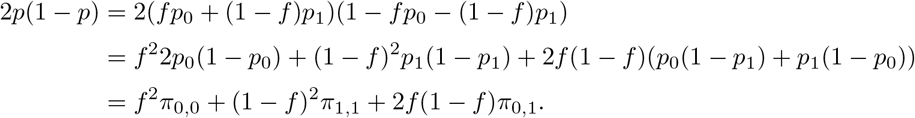

Here, *π*_*i,j*_ is the expected pairwise diversity between two samples, one drawn from population *i* and the other from population *j*. If *i* = *j*, this is pairwise diversity within a single population.

Plugging into the definition for *V*_*G*_, we get after admixture

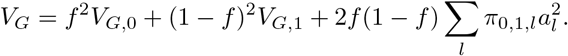

We can write *π*_0,1_ at a given locus in terms of *π*_0,0_, *π*_1,1_, and *F*_2_(0, 1) = (*p*_0_ − *p*_1_)^2^, a measure of single-locus allele frequency differentiation (Peter, 2016). Then

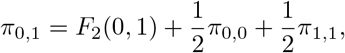

and

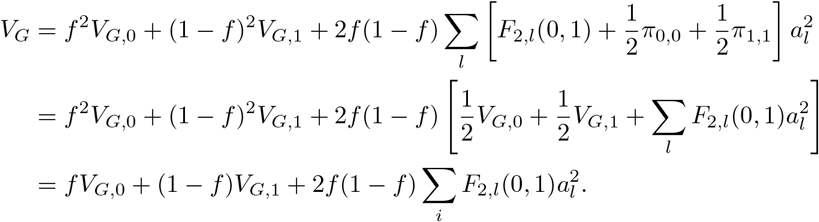

