## Supporting Information for "Archaic introgression and the distribution of shared variation under stabilizing selection"

### Supporting Information for
### "Archaic introgression and the distribution of shared variation under stabilizing selection"

Aaron P. Ragsdale

Department of Integrative Biology, University of Wisconsin–Madison, WI, USA

August 20, 2024

#### S1 Supplemental Figures

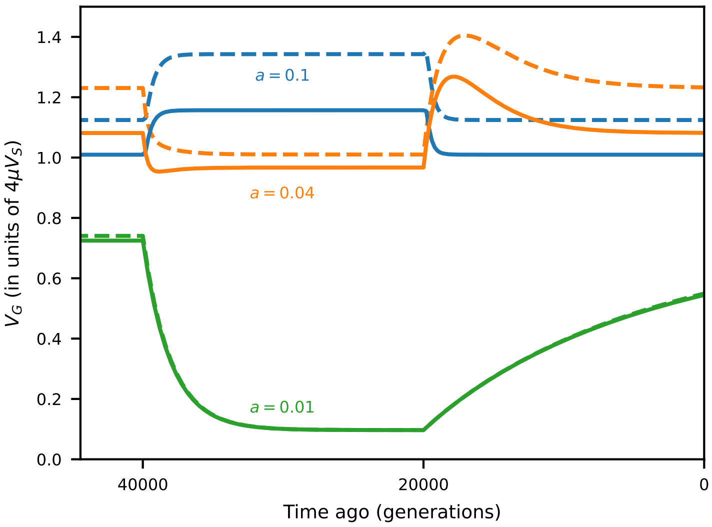


Figure S1: Single-population dynamics when $V_G$ is not small. Solid lines: expected $V_G$ assuming $s = a^2/2V_S$. Dashed lines: expected $V_G$ assuming $s = a^2/2(V_S + V_G)$. When mutation rates are large (here, $\mu = 0.01$, $V_S = 1$), $V_G$ is non-negligible compared to $V_S$. By having to account for $V_G$ in the translation of effect size to selection coefficient, it makes $s$ non-constant if $V_G$ changes over time (due to non-constant demography, for example). In this case, $s$ must be updated regularly, given the current state of the population. Here, the demographic history is a bottleneck followed by a recovery, as depicted in Figure 2A.

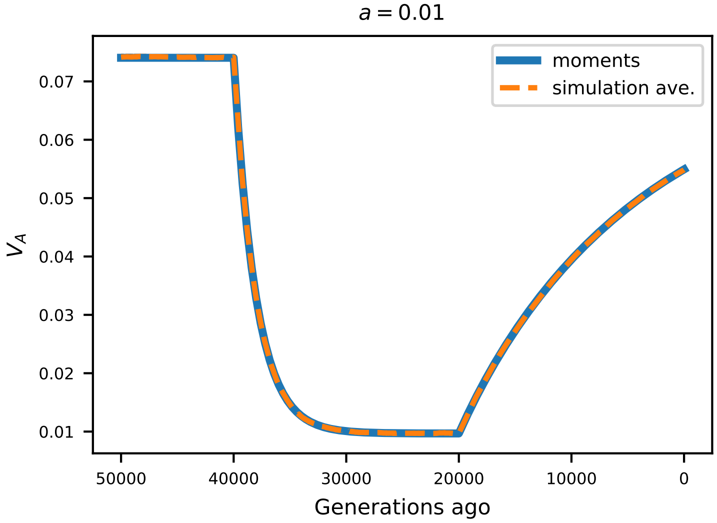


Figure S2: Simulations with no linkage and weak effects. Total mutation rate was $\mu = 0.025$ and all mutations had effect sizes $a = \pm 0.01$. The demographic history (bottleneck and recovery) is depicted in Figure 2A. The predicted trajectory of additive genetic variance using **moments** was found assuming symmetric underdominant selection on trait-affecting alleles, with selection coefficient $s = a^2/2(V_S + V_G)$.

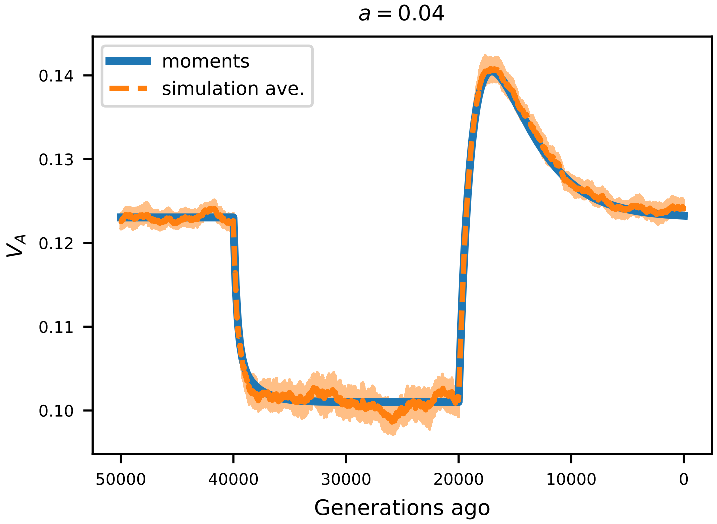


Figure S3: Simulations with no linkage and moderate effects. All parameters were consistent with Figure S2, but with effect sizes $a = \pm 0.04$.

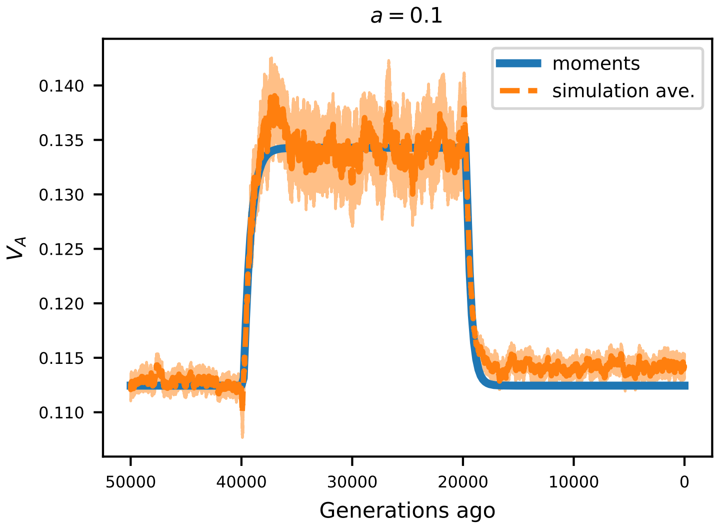


Figure S4: Simulations with no linkage and strong effects. All parameters were consistent with Figure S2, but with effect sizes $a = \pm 0.1$. In each comparison (weak, moderate, and strong effect sizes), predictions from `moments` closely match observed average additive genetic variance from simulations.

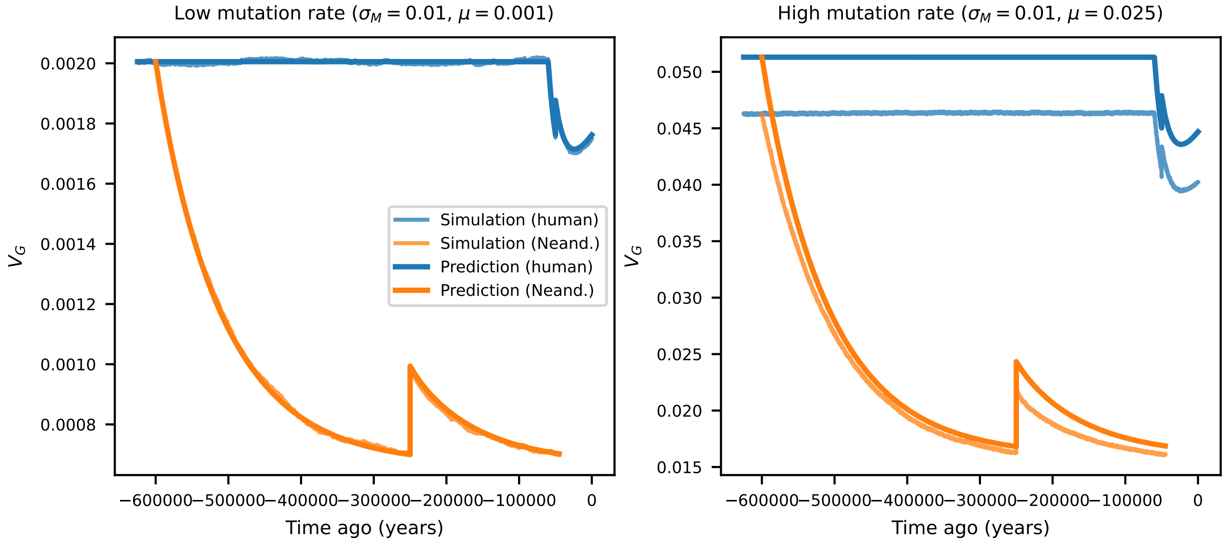


Figure S5: With low mutational variance ($V_M = 0.0001$), individual-based simulations using `fwdpy11` closely match our predictions using `moments`, when the mutation rate is small (Left). When the mutation rate is increased, polygenicity increases and unlinked expectations from `moments` deviate from observed $V_G$ in individual-based simulations. In this case, with relatively small mutational variance, the observed $V_G$ is reduced in comparison, consistent with the Bulmber (1971) effect.

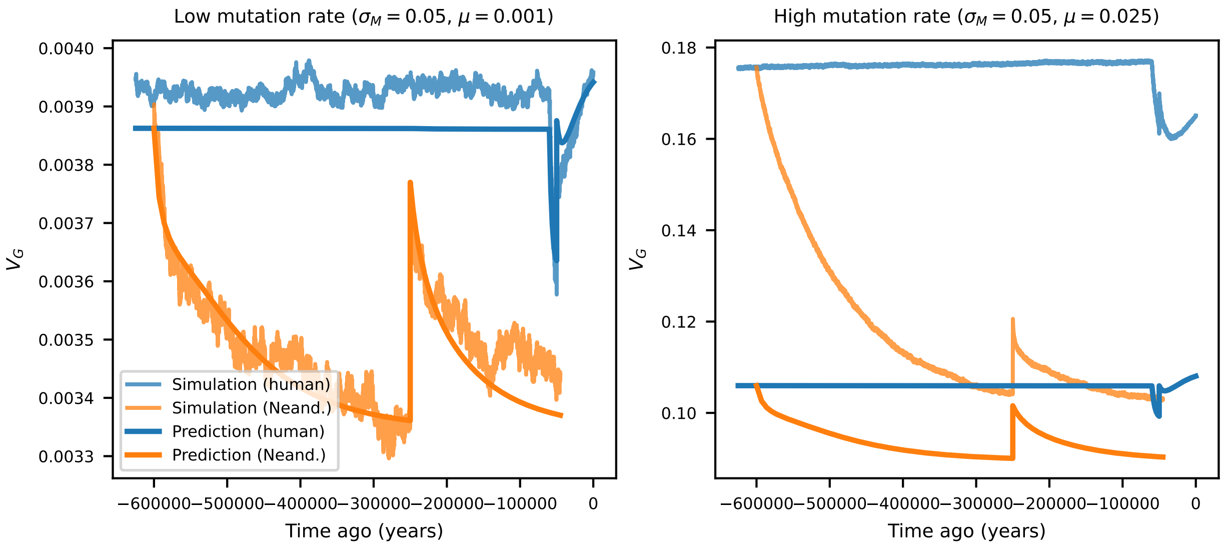


Figure S6: With high mutational variance ($V_M = 0.0025$), strong deviations are observed between individual-based simulations and expectations without linkage. When the mutational input is large, observed $V_G$ can be much larger than expectations without linkage, possibly consistent with widespread interference between alleles.

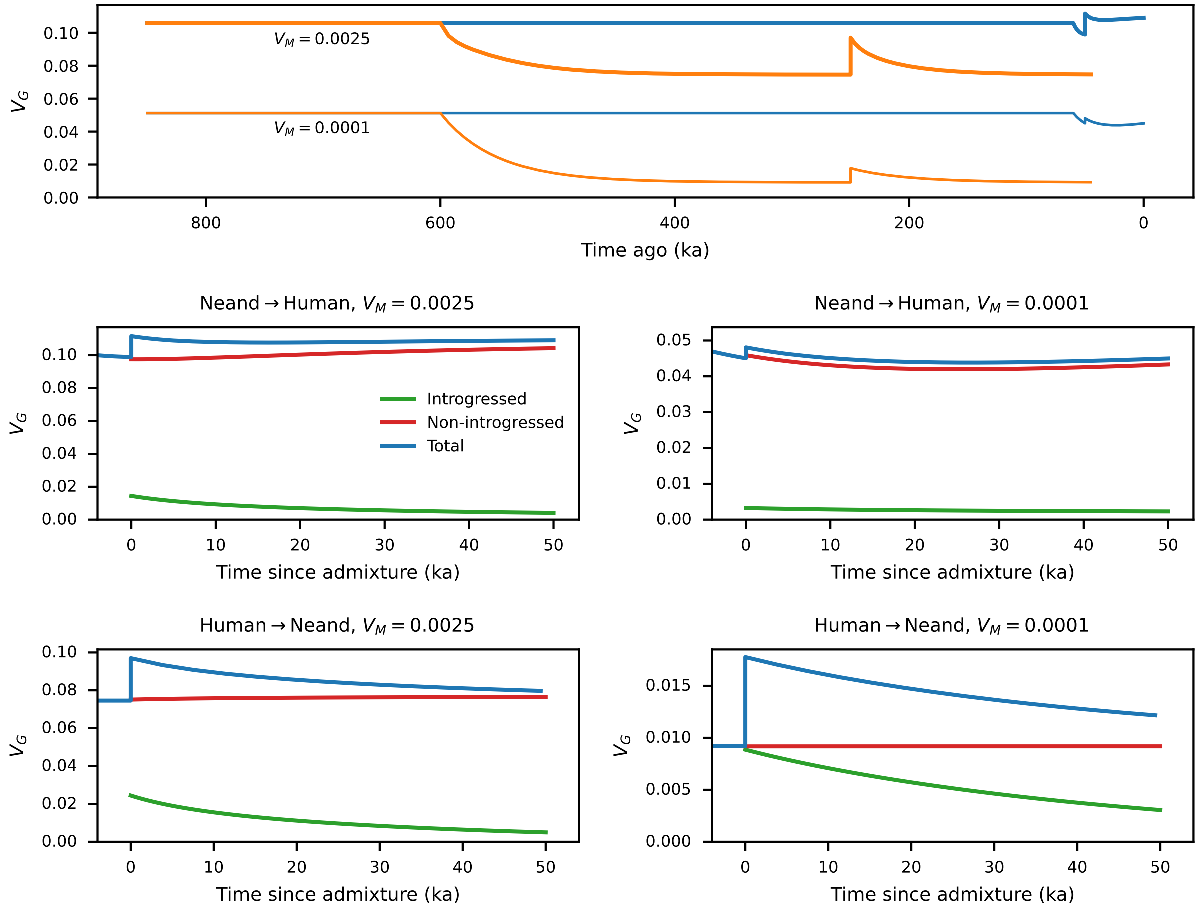


Figure S7: Genetic variance of a trait under stabilizing selection in a model with human (blue) and Neanderthal (orange) demography. We consider two mutational variances ($\sigma_M = 0.05$ and $0.01$). The demographic model (Figure 4A) includes 5% admixture from humans to Neanderthals 250 ka, and 2% admixture from Neanderthals to humans 50 ka.

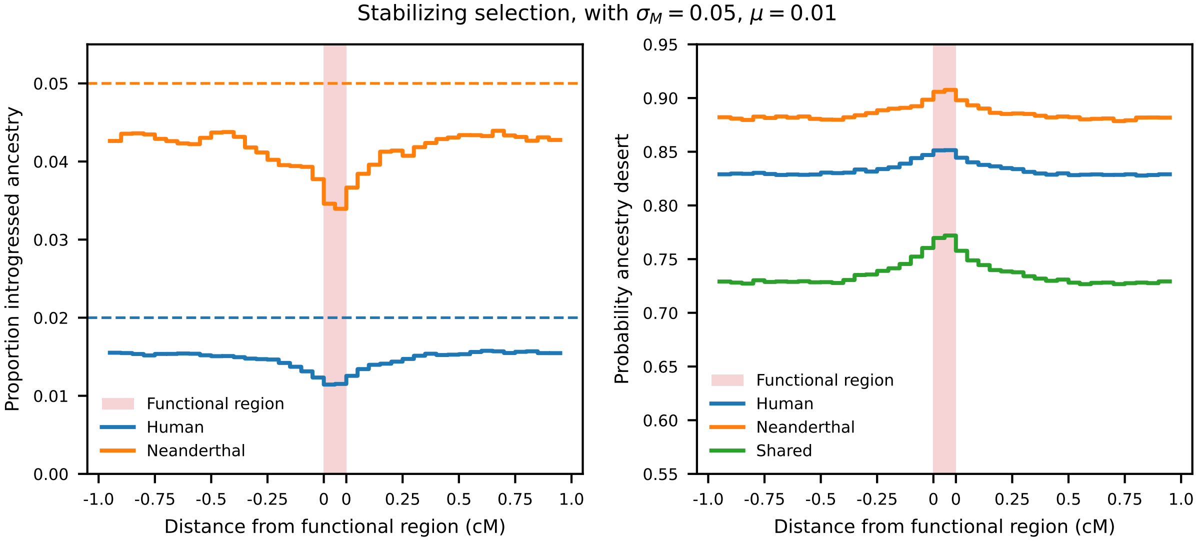


Figure S8: Introgressed ancestry and probability of introgression deserts in and around functional regions, from simulations with high mutational variance. The demographic model is shown in Figure 4A.

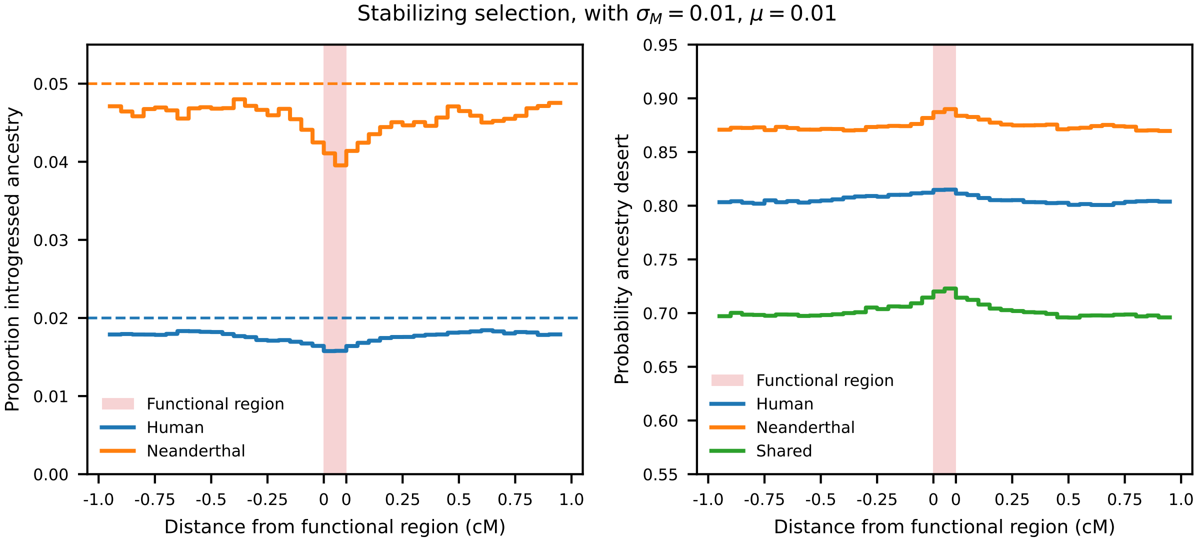


Figure S9: Introgressed ancestry and probability of introgression deserts in and around functional regions, from simulations with moderate mutational variance. The demographic model is shown in Figure 4A.

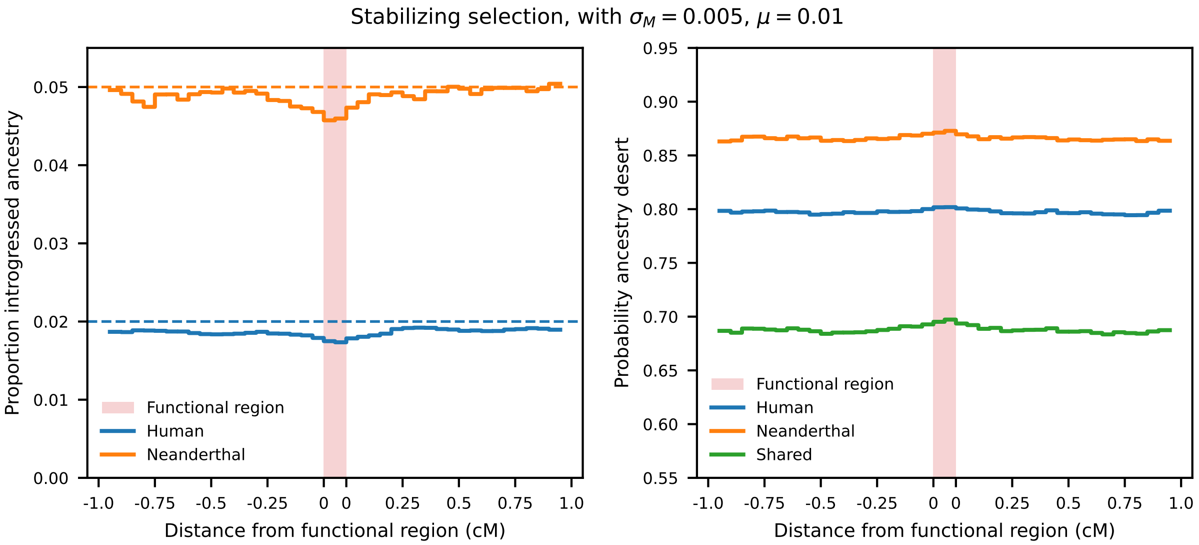


Figure S10: Introgressed ancestry and probability of introgression deserts in and around functional regions, from simulations with low mutational variance. The demographic model is shown in Figure 4A.

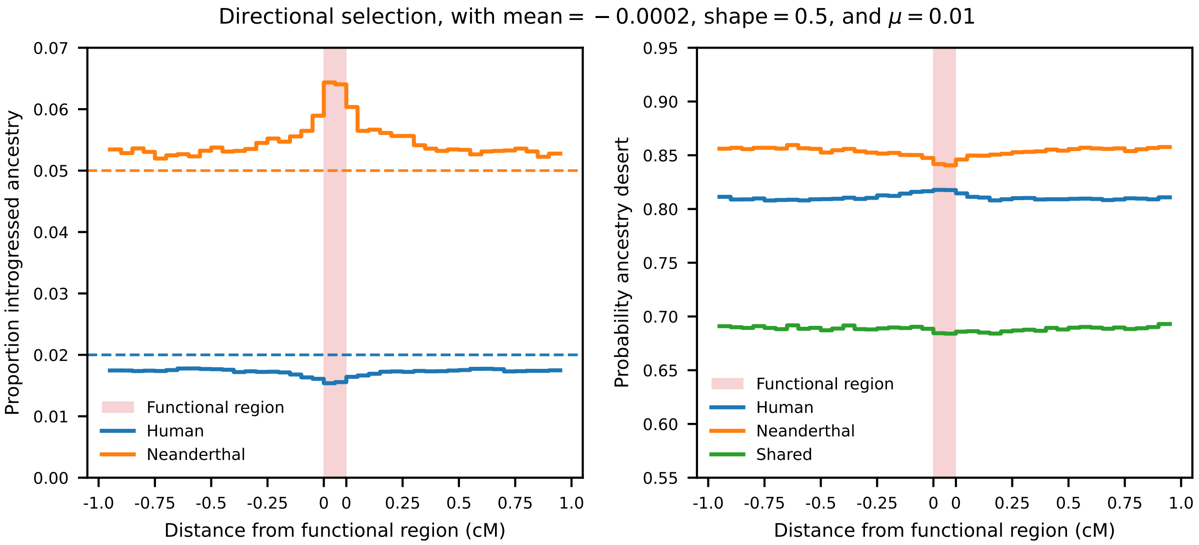


Figure S11: Introgressed ancestry and probability of introgression deserts in and around functional regions, from simulations with directional selection. Deleterious mutations were drawn from a gamma distribution with mean $-0.0002$ and shape 0.05. The demographic model is shown in Figure 4A.

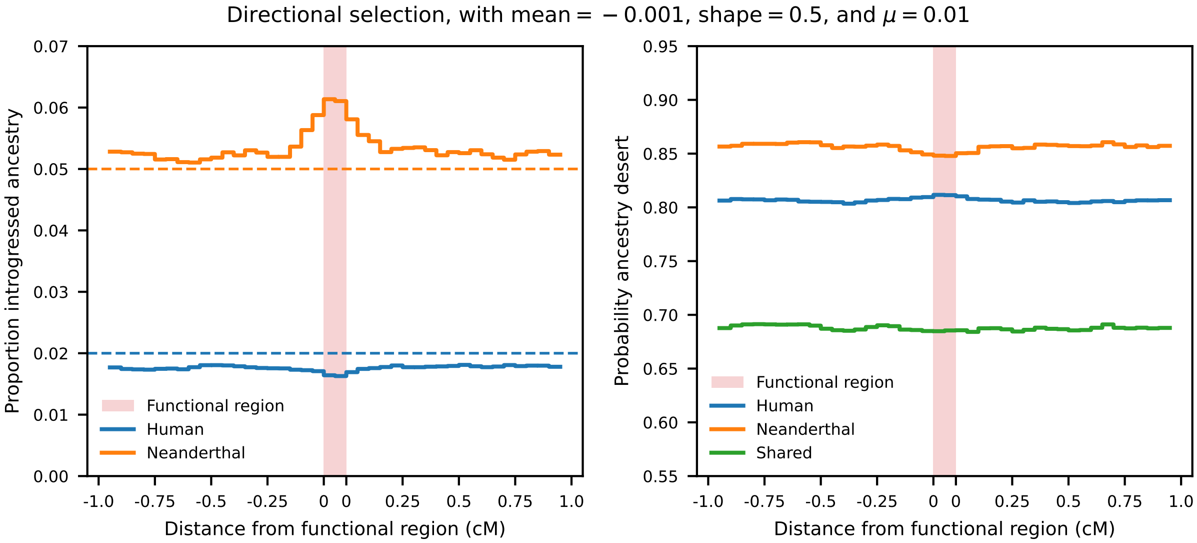


Figure S12: Introgressed ancestry and probability of introgression deserts in and around functional regions, from simulations with directional selection. Deleterious mutations were drawn from a gamma distribution with mean $-0.001$ and shape 0.05. The demographic model is shown in Figure 4A.

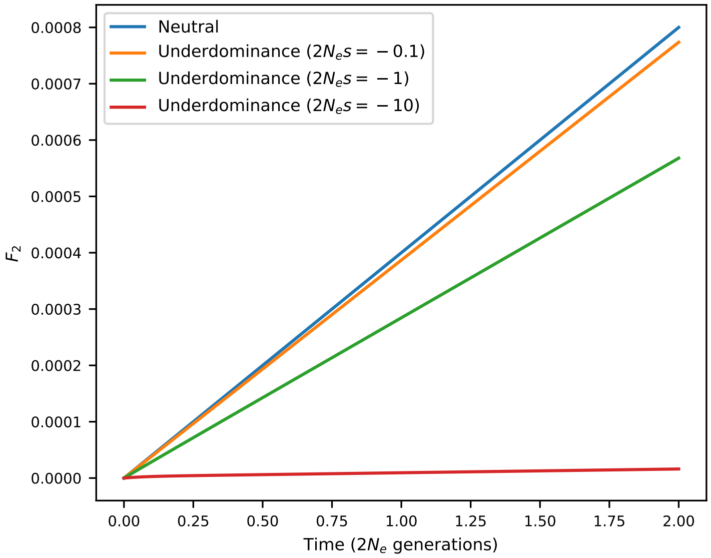


Figure S13: Underdominance, like negative selection, reduces expected $F_2$ compared to neutral divergence.

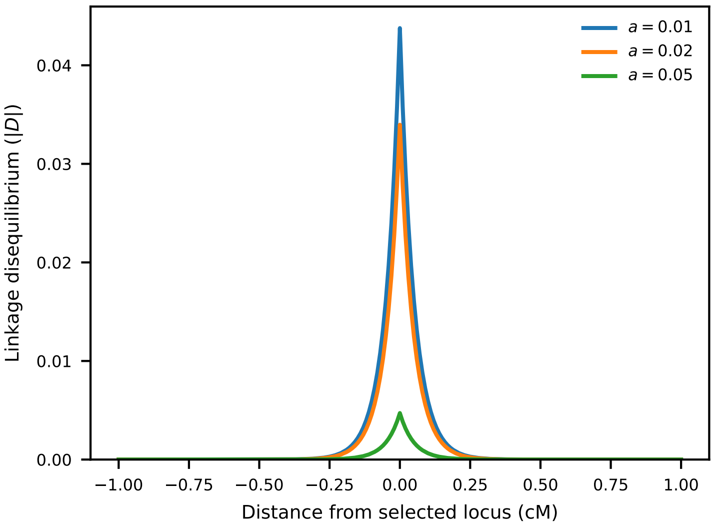


Figure S14: Linkage disequilibrium between the a trait-affecting and neutral allele, 2000 generations after admixture. Initially, the introgression proportion was 0.05. Mutations with stronger effect sizes, and thus stronger selection against them, decrease in frequency more rapidly, leading to reduced LD as measured by $D = Cov(p, q)$ (Figure 5).

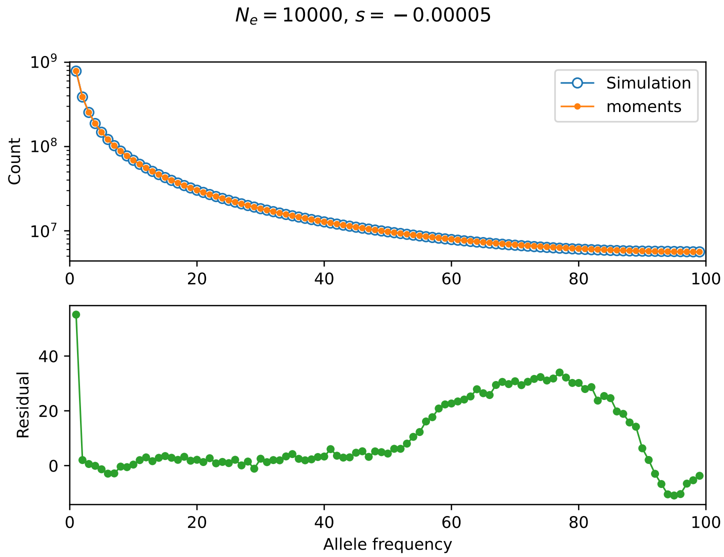


Figure S15: Comparison of predicted (`moments`) and simulated SFS with underdominant selection. In this comparison, $N_e = 10^4$ and $s = -5 \times 10^{-5}$, so that the population-size scale selection coefficient $\gamma = -1$. Simulations were performed under a Wright-Fisher model without linkage.

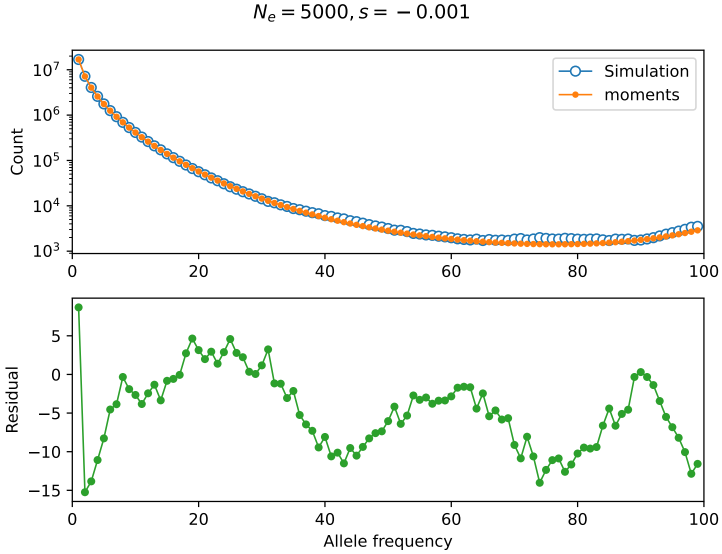


Figure S16: Comparison of predicted (`moments`) and simulated SFS with underdominant selection. In this comparison, $N_e = 5000$ and $s = -0.001$, so that the population-size scale selection coefficient $\gamma = -10$. Simulations were performed under a Wright-Fisher model without linkage.

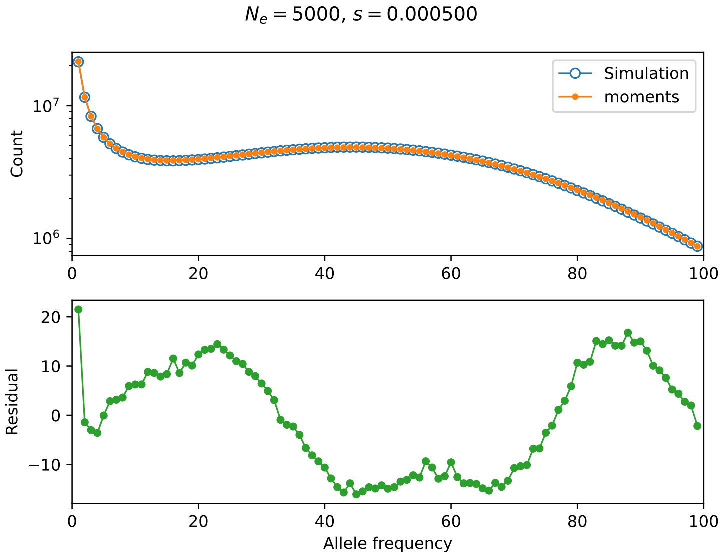


Figure S17: Comparison of predicted (`moments`) and simulated SFS with *over*dominant selection. In this comparison, $N_e = 5000$ and $s = 0.0005$, so that the population-size scale selection coefficient $\gamma = 5$. Thus, with positive selection, heterozygotes are favored. Simulations were performed under a Wright-Fisher model without linkage.
